# From microbiome to sperm motility traits: An inside out perspective

**DOI:** 10.1101/2025.11.28.691116

**Authors:** Marco Graziano, Gian Luigi Garbini, Alessandro Devigili, Livia Pinzoni, Andrea Quagliariello, Emanuele Bosi, Clelia Gasparini, Marco Fondi, Maria Elena Martino

## Abstract

Growing interest in the relationship between microbiome composition and host biology has revealed the many ways host-associated microbes influence physiology, ecology, and evolution. However, microbial communities associated with reproductive organs - and their roles in reproduction - remain poorly understood. Here, we characterized the skin- and ejaculate-associated microbiomes in an internally fertilizing fish and tested whether microbial diversity and specific bacterial taxa correlate with sperm motility traits key for reproductive success. We used the guppy (*Poecilia reticulata*), a well-established model in ecology and evolutionary biology with well-characterised reproductive physiology. In guppies, sperm velocity is a validated predictor of male reproductive performance, making them a powerful system for exploring microbiome–fertility interactions. Our analyses reveal a correlation between skin microbiome diversity and sperm performance. Notably, increased skin microbiome total richness is associated with reduced sperm velocity, whereas no significant associations were detected for ejaculate-associated microbiomes. We also identified bacterial taxa across both tissues that were positively or negatively linked with sperm performance. These findings suggest that, while the ejaculate-associated microbiome may directly influence sperm traits, the skin microbiome could serve as a proxy for reproductive potential by reflecting systemic physiological and immunological states associated with fertility.

## Introduction

Animal microbiomes are increasingly recognised as integral components of host biology, with the hologenome theory emphasizing their evolutionary and ecological significance [1–4]. While extensive research has explored the roles of gut, oral, and skin microbiomes, reproductive microbiomes (RMs) remain understudied. Yet, given that reproduction is the primary target of natural and sexual selection, overlooking the role of microbiomes in this context leaves a critical gap in our understanding of organismal biology [5–7].

Over the last three decades, research on RMs has focused mainly on humans [8,9] and invertebrates, particularly those associated with the iconic Alphaproteobacteria of the genus *Wolbachia* [10,11]. While much is known about pathogenic effects on reproduction [12–16], the broader ecological role of microbiomes in fertility are poorly understood.

In addition, host–microbiome interactions extend across body compartments [17], and, while RMs may directly affect fertility, other microbiomes, like the gut and skin, may also influence or predict reproductive potential. Because host-associated microbial communities are tightly linked to immune function, endocrine activity, and overall physiological state, external microbiomes could serve as indicators of internal health, including reproductive condition [18,19]. The skin, as the primary interface with the environment and external microbes, has been proposed as an indicator of overall health [20]. In this context, variation in the skin microbiome may mirror systemic physiological differences among males, thus providing an externally accessible signal of reproductive capability.

This study aims to investigate how skin- and ejaculate-associated microbiota relate to sperm motility traits using guppies (*Poecilia reticulata*), a well-established model in reproductive research [21–25]. Guppies are internally fertilizing livebearers: males possess a modified anal fin (gonopodium) that functions as an intromittent organ to transfer sperm directly into the female reproductive tract. This, distinguishes guppies from most externally fertilizing fishes, providing a unique opportunity to examine host–microbe interactions in reproductive tissues. Because fertilization occurs within the female, the influence of external environmental microbes is minimized, allowing greater experimental control and clearer inference of host-driven microbial and reproductive dynamics. Internal fertilization requires strict immunological regulation of the sperm environment, where microbial colonization may occur within ejaculates and associated tissues but is highly shaped by host processes rather than ambient exposure [26–28]. Understanding how these localized microbiomes relate to sperm performance can therefore reveal how host–microbe dynamics influence reproductive potential in natural mating systems. Here, using sperm velocity as a proxy for fertilisation success [29], we assessed whether microbial diversity and the relative abundance of specific taxa in skin and sperm-associated microbiotas correlate with key sperm motility traits known to influence reproductive success.

## Methods

### Fish origin and husbandry

Guppies (*Poecilia reticulata*) used in this study were descendants of wild-caught individuals (Lower Tacarigua River, Trinidad, 2002). Since 2013, progeny have been maintained in semi-natural conditions in a pond at the University of Padova. In the lab, fish were housed in 150 L tanks (n ≈ 50, 1:1 sex ratio) at 26 ± 1°C (12:12 h light/dark). Fish received a mixed diet of *Artemia salina* nauplii and dry food (TetraMin®), with weekly Chironomid larvae. During maintenance, individuals were rotated among tanks to ensure adequate outbreeding and homogenize environmental variation. When new tanks were established, they were founded with fry originating from approximately 10 different source tanks, produced by ∼30 mothers and sired by an estimated ∼90 fathers, given the promiscuous mating system of guppies. (see [30]).

### Microbiota isolation

Fully mature males (n= 16) of the same age cohort (6 months old) were randomly selected and individually anaesthetized in a beaker containing their own system water (i.e., water drawn directly from the holding tank) supplemented with 0.02% tricaine methanesulfonate (MS-222). Each fish was briefly rinsed with sterile water, placed onto a sterile Petri-dish and swabbed ten times with a sterile swab, from behind the operculum to the caudal peduncle. The swab was placed in a sterile 2.5 ml Eppendorf containing 1 mL of RNAlater® solution. Sperm were collected by gently applying pressure to the male’s abdomen, causing the release of sperm in the Petri dish containing physiological solution (0.9 % NaCl). In guppies, sperm are released as sperm bundles, which are discrete aggregations of tightly packed spermatozoa (spermatozeugmata). Half of the collected ejaculate sample was immediately processed for sperm traits analysis, whereas the remaining half was placed in RNAlater® as described for the skin swabs (figure S1) and frozen at -20 °C until RNA extraction. All sampling and RNA extractions were carried out using aseptic procedures. Comprehensive contamination-control are detailed in the Supplementary Material (SP1).

### RNA extraction and 16S rRNA sequencing

RNA was extracted using the RNAeasy Mini Kit (Qiagen) following the manufacturer’s protocol, then reverse-transcribed to cDNA (SuperScript™ IV, Thermofisher). Amplicon sequencing of the bacterial 16S rRNA gene targeted the V3–V4 region using the universal primers 341F and 806R. Library preparation and sequencing were outsourced (BMKGENE, Germany) on an Illumina GAiiX as paired-end reads (PE250). Sequences were processed in QIIME2[31]. Denoising and chimera removal used DADA2[32]; singleton and doubleton were also removed from the dataset and the data was rarefied at a depth of 105.000. Taxonomy was assigned using a naïve Bayes classifier trained on SILVA (v. 138.2, https://silva.arb.de) as described in [33],[34].

### Bioinformatics

The amplicon sequence variant (ASV) abundance table from the denoising, the taxonomy table, and the file with the experimental metadata were merged in a phyloseq object [35] in R (4.2.2). To better highlight differences in bacterial abundances across samples and conditions, the dataset was normalized using z-scores, which transform each value based on its deviation from the mean. The resulting standardized values were then used to generate the heatmap. A rooted phylogenetic tree was constructed using the align-to-tree-mafft-fasttree pipeline from the q2-phylogeny plugin [31,36]. This process included multiple sequence alignment via MAFFT, alignment masking, and tree inference using FastTree, enabling computation of phylogenetic diversity metrics and UniFrac distances.

Alpha diversity metrics (Chao1 and Shannon) were calculated using the *phyloseq* package. Statistical differences between skin and ejaculate samples were tested using non-parametric ANOVA (Wilcoxon tests). Beta diversity was calculated using weighted UniFrac distances. Taxa were summarized at the genus level where possible. The term ‘Unclassified’ refers to sequences confidently assigned to a higher taxonomic rank but not to any named genus in the reference database. ‘Other’ denotes a residual category aggregating very low-abundance genera (each <0.1% mean relative abundance). Sample ordination was performed via Principal Coordinate Analysis (PCoA), and group differences were evaluated using PERMANOVA [37] via the adonis2 function (*vegan* package [38]).

Association networks were built using Spearman correlations. Only significant correlations were included (p< 0.01) with thresholds either < -0.85 or > 0.85. Within the networks, each node represents the genus and each edge the significant pairwise correlation between them. Positive correlations between two genera denoted similar abundance patterns, while negative ones were characterised by opposite abundance patterns. Interacting nodes within networks represented co-occurrence across samples. After networks construction, topological features were measured using Cytoscape 3.10.3 [39] and are detailed in the Supplementary Material (SP2).

### Sperm motility traits assessment

Sperm traits were assessed following established CASA protocols [40,41]. Briefly, sperm were activated with a 150 mM KCl + 2 mg mL:::¹ BSA solution and analysed using a CEROS computer-assisted sperm analyser (Hamilton-Thorne) coupled with a Nikon Eclipse E200 phase-contrast microscope. Sperm performance analysis included curvilinear velocity (VCL, μm/s), average path velocity (VAP, μm/s), straight line velocity (VSL, μm/s), linearity (LIN, %), beat cross frequency (BCF, Hz), straightness (STR) and amplitude of lateral head displacement (ALH). Further details about CASA acquisition settings are summarized in the Supplementary table S1. For each male, motility was independently assessed in three distinct sperm-bundle groups obtained from the **s**ame ejaculate, with an average of 68.23 ± 18.64 SD motile sperm cells analysed per each replicate. Motility traits were checked for distributional properties, repeatability across replicates, and inter-trait correlations; highly repeatable and collinear traits were subsequently summarised via PCA. Because microbiome diversity was measured once per tissue per male, and to avoid pseudoreplication, motility traits were averaged across the three replicates prior to all microbiome–trait modelling.

A full description of repeatability analyses, distributional checks, PCA methodology, and loading tables is provided in the Supplementary Material (SP3).

### Trait–Richness Models

For our response variable, VCL (and also separately for PC1 and PC2 whose results are reported in the supplementary material), we applied an automated, multi-step model-selection pipeline (R v4.4.2) that included both Chao1 richness and Shannon entropy as alternative diversity predictors. Models were ranked using AICc, and only diversity metrics supported by this automated filtering procedure were retained. To test whether sperm performance covaried with microbial diversity, we modelled sperm velocity (VCL) as a function of Chao1/Shannon and tissue identity (skin vs. sperm bundles). All models incorporated the interaction between diversity and tissue identity, and raw and z-standardised predictors were evaluated. We fitted a comprehensive grid of candidate models including linear models (LMs), linear mixed-effects models (LMMs), and generalized linear mixed-effects models (GLMMs, via *glmmTMB*), comparing structures with and without a random intercept for male identity to account for paired observations (each male contributing one skin and one sperm-bundle sample). Alternative model formulations, random-effect structures, and scaling options were compared using AICc, and the best-supported model for each response was selected from this full candidate set.

In parallel, we evaluated tissue-specific effects by fitting models where microbial diversity was entered as separate compartment-specific predictors (e.g., Chao1_skin and Chao1_bundle). Model assumptions and residual structure were verified with DHARMa. You can find indicated supplementary material in the dedicated section, present.

Details on model-selection workflow, diagnostics, and the equivalent analyses for PC1 and PC2, is provided in the Supplementary Material (SP4).

### Multivariate Analyses of Trait–Microbiome Relationships

To assess how variation in microbial community composition related to sperm traits, we used constrained ordination (CCA and distance-based RDA). A two-stage workflow was applied: (1) automated selection of filtering thresholds and data transformations, and (2) final pooled and within-tissue models relating community structure to either VCL or PC1 + PC2. Permutation tests were blocked by fish ID. Key microbial genera were identified from species scores and vector alignment.

Comprehensive methodological details, parameter sweeps, diagnostics, and taxa loading statistics are reported in the Supplementary Material (SP5).

## Results

We began by profiling the microbiota associated with the skin and the ejaculate of 16 male guppies (a complete list of detected ASVs is provided in the Electronic Supplementary Material: *“Genus_ab_165.txt”*). After quality control, microbial data were available for all sperm-bundle samples and for 15 of 16 skin samples, as one skin sample showed RNA degradation and was therefore excluded from further analyses. Alpha diversity analysis revealed that skin-associated microbiomes exhibit significantly higher bacterial total richness (Chao1 index) compared to ejaculate-associated microbiomes. This difference reflects the skin’s richer pool of low-abundance ASVs, rather than broader taxonomic breadth, as both tissues contained an equivalent number of detected genera (164 for the skin and 165 for the ejaculates, exceeding the 0.25% relative-abundance threshold. Shannon index did not show a statistically significant difference, although a trend toward higher diversity in skin microbiomes was detected (figure 1A). Beta diversity analyses showed significantly distinct clustering of microbial communities associated to skin and ejaculates (PERMANOVA, p <0.01, R^2^= 0.94, figure 1B), with some shared taxa. Ejaculate samples were richer in *Massilia*, *Acinetobacter*, *Lactobacillales*, and *Brevundimonas*, while *Prevotellaceae*, *Clostridia*, and *Rikenellaceae* dominated in skin (Figure 1C, D, E, Supplementary figure S3, Table S2). *Fluvicola* was uniquely found in ejaculate. Notably, both tissues show considerable inter-individual variation in the relative abundance of these genera (Figure 1D, E). This variability is often due to the high abundance of specific genera, such as *Mycobacterium* (e.g., SS_13, SS_14, B13).

**Figure 1.**
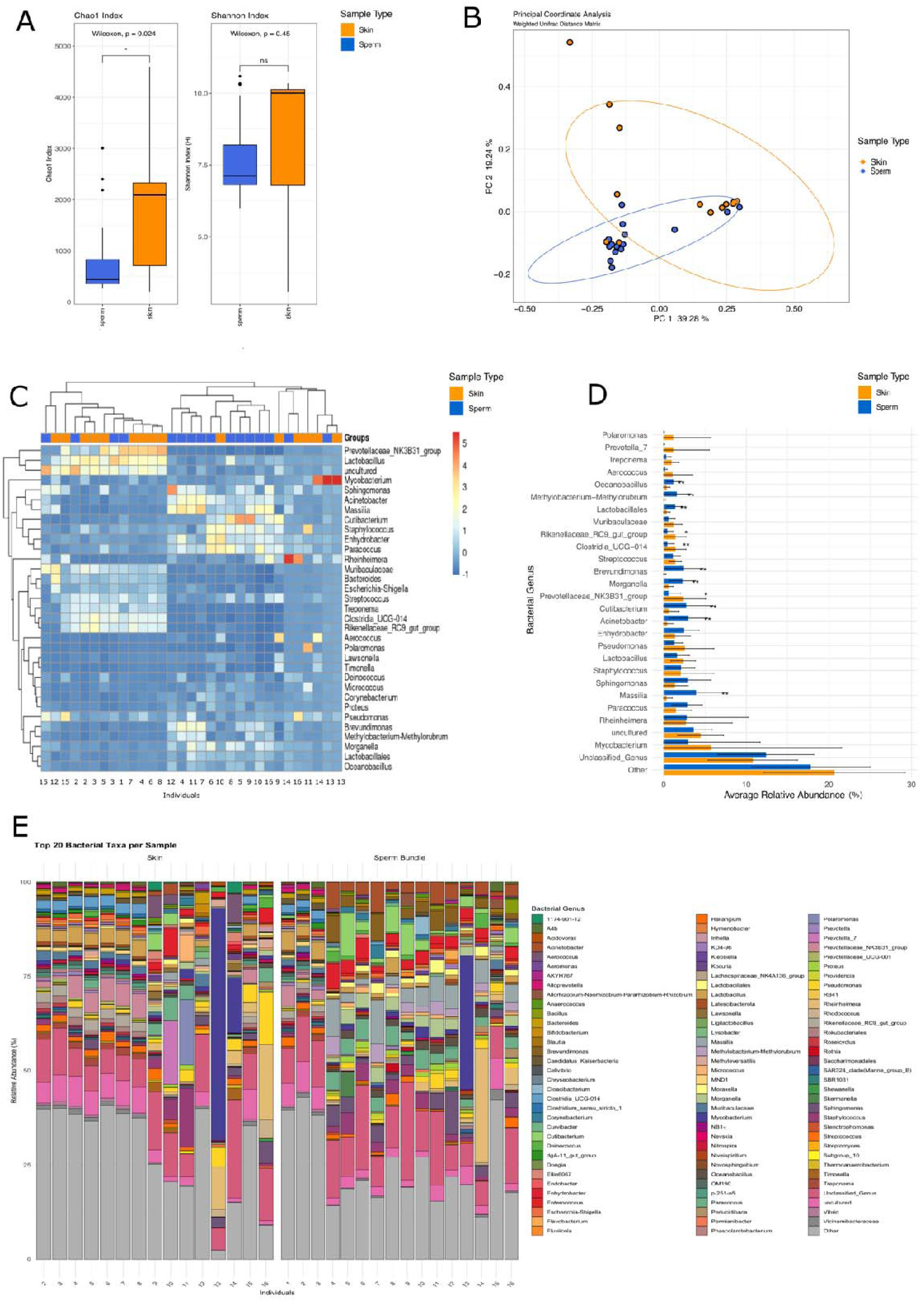
**Alpha (A) and Beta diversity (B) indices** for sperm bundles (blue) and skin swabs (orange); **(C)** Heatmap showing clustering and z-scores of bacterial abundances in the two sample types; **(D) Barplot** reporting the twenty most abundant bacterial taxa by sample type. Error bars represent the standard deviation across 16 individuals for each taxon. Asterisks indicate statistically significant differences between tissues (* = p<LJ0.05, ** = p<LJ0.01, *** = p<LJ0.001); **(E) Relative abundances** of bacterial taxa higher than 0.25% per sample type.

### Principal component structure of sperm motility traits

We first examined the structure of sperm motility variation using principal component analysis (PCA; Supplementary Figure S2). The first two components (PC1 and PC2) captured 87.3% of total variance, highlighting two distinct functional dimensions in sperm kinematic traits. PC1 was associated with sperm velocity parameters (VAP, VSL, VCL), whereas PC2 represented variation in directionality measures (ALH, LIN, STR). Because VCL predicts reproductive success in guppies [29], subsequent analyses focused primarily on this parameter as a proxy for overall sperm performance. Nevertheless, as the overarching aim of this project is to describe the relationship between sperm traits and microbiome composition, we do report results including all the sperm motility parameters based on PCA analysis (table S4 and S5).

### Modelling sperm velocity as a function of microbial richness

To test whether sperm velocity (VCL) covaried with microbial diversity derived from either the skin or the sperm bundles of the same males, both Chao1 richness and Shannon entropy were initially included as candidate diversity metrics; however, only Chao1 was consistently retained by AICc-based filtering. To test for compartment-specific effects, we first fitted interaction models of the form *VCL ∼ Chao1 × Tissue*. This model identified a significant negative association between microbial richness and sperm velocity (β = –0.0089 ± 0.0036 SE, p = 0.021; Table S4), such that males with more diverse microbiota tended to exhibit slower sperm. Neither the tissue main effect nor the interaction term were significant, indicating that the negative slope was similar across tissues. Equivalent models using PC1 and PC2 scores showed qualitatively similar but weaker trends (Table S4).

Because our biological question concerned whether diversity from each compartment independently predicted sperm performance—rather than the effect of tissue identity per se—we next fitted models with separate predictors for each compartment (*VCL ∼ ChaoSkin + ChaoBundles*). This formulation avoids pseudoreplication and allows direct assessment of tissue-specific associations. In this model, skin microbial richness showed a strong negative effect on sperm velocity (β = –10.42, p = 0.002; Figure 2A; Table S5), whereas ejaculate richness was not significant. Parallel analyses using PC1 and PC2 supported the same overall pattern and are reported in the Supplementary Material (Table S4, S5).

**Figure 2.**
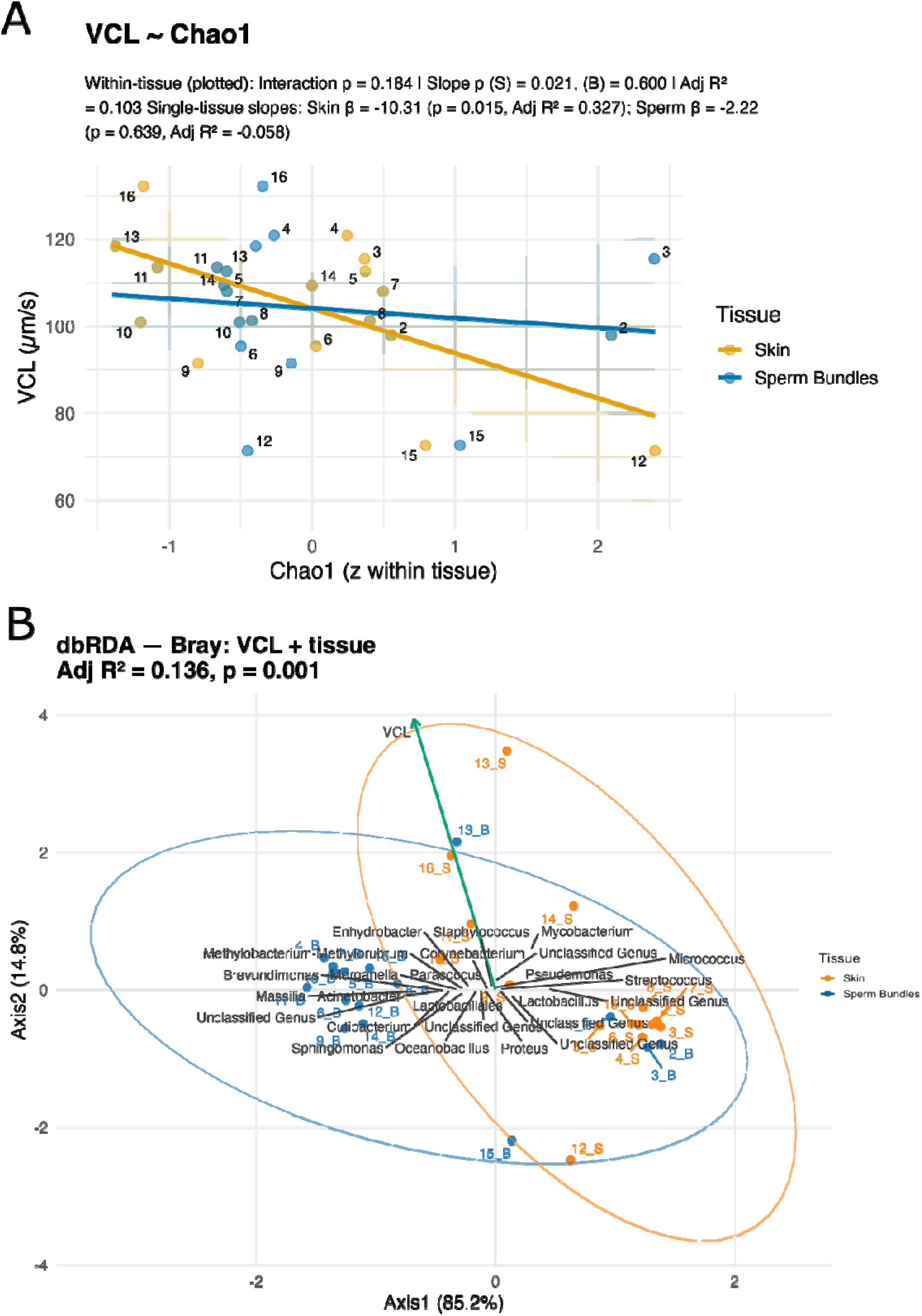
**(A) Relationship between sperm velocity (VCL) and standardized bacterial richness (Chao1)** within skin (orange) and sperm bundles (blue). Each point represents an individual male (numbers correspond to fish IDs). Lines show fitted linear relationships (±95% confidence intervals). **(B) Distance-based redundancy analysis (dbRDA, Bray–Curtis) showing bacterial community composition constrained by VCL and tissue type**. Ellipses (95% confidence) illustrate tissue-level clustering. The VCL vector indicates the direction of increasing sperm velocity and its association with specific genera.

### Multivariate associations between sperm traits and microbiome composition

The automated model sweep revealed consistent effects of filtering on model performance. Both CCA and dbRDA models improved markedly under moderate prevalence and abundance thresholds, with dbRDA explaining the greatest proportion of variance across all tested configurations. The selected parameters (prevalence ≥ 5; mean RA ≥ 0.005) yielded reproducible and well-constrained ordinations.

In the optimized models using the top 25 genera, VCL and sperm trait principal components were strongly associated with microbial community composition when tissue identity was included as a covariate. The best-performing models explained up to 14.5% of the constrained variance for PC1 + PC2 and 13.3% for VCL (both p < 0.001), Figure 2B supplementary Figure S3, electronic supplementary files (Dryad)), compared with 8–10% in the corresponding CCA analyses. Including tissue as a predictor markedly improved fit, increasing adjusted R² from approximately 0.07 (trait-only models) to 0.13–0.15, indicating that tissue identity alone accounted from one third to one half of the total explained variance. In ordination space, skin and sperm-bundle samples formed two clearly separated clusters, confirming strong tissue-specific structuring. The sperm traits explained a smaller but significant fraction of residual variation, suggesting that differences in sperm performance covary with microbial community composition across tissues (Figure 2B and S3). When tissue was partialled out from the models, the explained variance dropped sharply, and no within-tissue models reached statistical significance (adjusted R² ≈ 0.02; p > 0.9). Analyses performed separately for sperm bundles and skin samples confirmed the absence of significant relationships between single genera and sperm traits within compartment (adjusted R² ≤ 0.05; p > 0.3). These results indicate that the associations between sperm traits and microbiome composition are largely driven by between-tissue differences rather than by independent variation within tissues. In other words, sperm phenotype and microbial structure covary because both reflect tissue-specific physiological environments, not direct within-tissue coupling.

### Microbial taxa influencing ordination structure

The ordination biplots and species score analyses identified a subset of bacterial genera that most strongly aligned with the sperm trait axes (Tables S6 and S7), irrespective of their compartment of origin In particular, *Mycobacterium*, *Massilia*, *Acinetobacter*, and *Brevundimonas* exhibited the strongest vector lengths, indicating major contributions to VCL-associated community structure. *Mycobacterium* and *Pseudomonas* vectors showed strong positive alignment with increasing VCL, indicating that these genera were more abundant in individuals or tissues associated with higher sperm motility. Conversely, *Lactobacillus* and Lactobacillales vectors were oriented in the opposite direction, suggesting reduced prevalence under similar conditions. Intermediate taxa such as *Brevundimonas*, *Methylobacterium*, *Massilia*, and *Acinetobacter* showed moderate associations along the composite sperm trait axes. These genera significantly contributed to the overall structure of the ordination, suggesting that multiple bacterial lineages covary with different aspects of sperm swimming performance. Full dbRDA and CCA statistics, including model selection thresholds and corresponding adjusted R² values, are provided in electronic supplementary material.

To further investigate interactions among bacterial taxa, co-occurrence networks were built for each tissue (Figure 3). The skin microbiome showed a denser network (95 nodes, 276 edges, 4 modules) than the sperm bundle microbiome (62 nodes, 138 edges, 3 modules), indicating higher number of microbial interactions in the skin. Despite structural differences, both networks had similar modularity (skin: 0.446; sperm: 0.487, Figure 3B-D, Table S3), reflecting comparable compartmentalization. Key hub taxa identified through betweenness centrality included *Lactobacillus* and *Vibrio* in the skin, *Candidatus Solidibacter* in the ejaculate-associated microbiome, and *Prevotellaceae* in both compartments.

**Figure 3.**
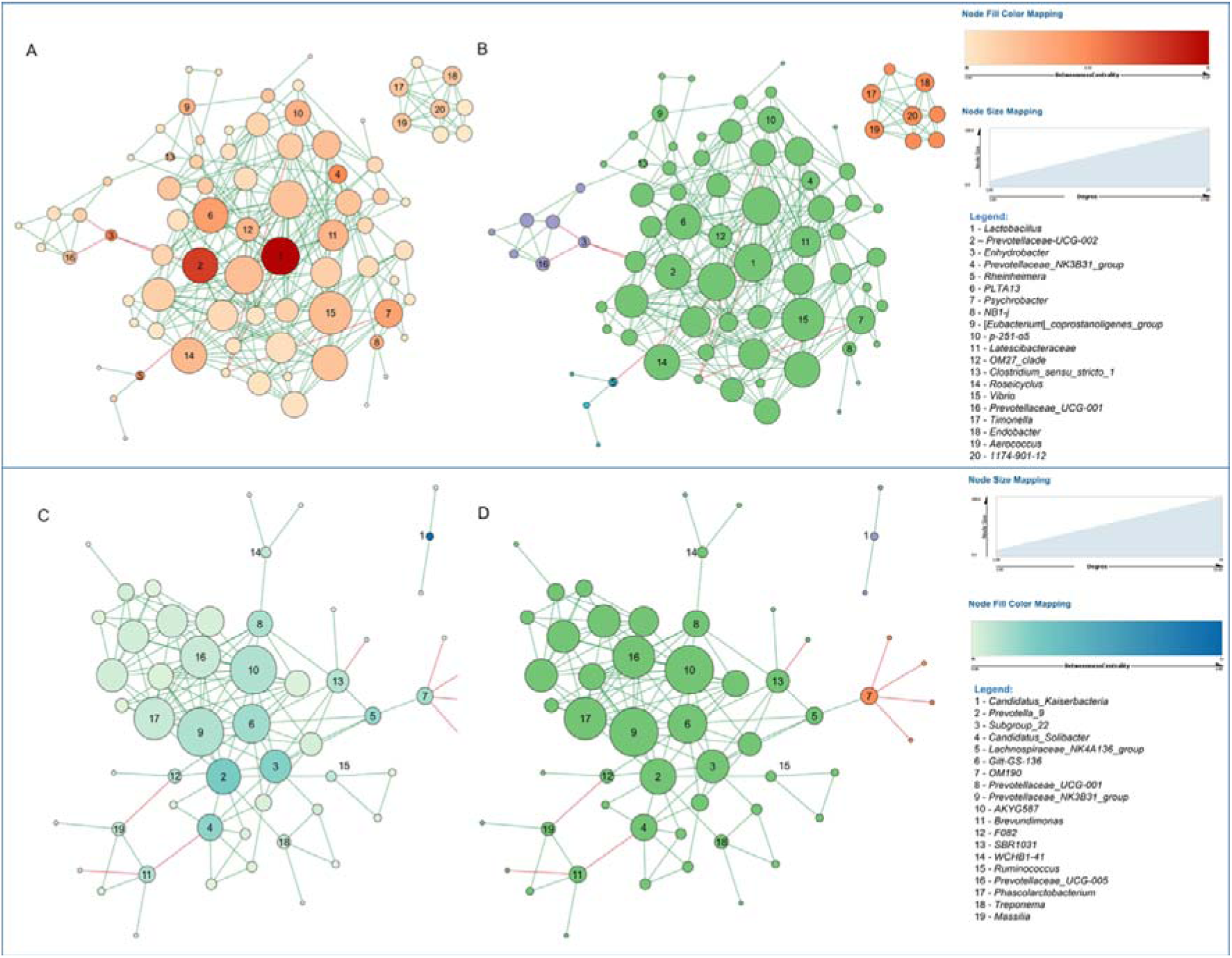
Visualization of bacterial networks for skin. **(A-B)** and sperm **(C-D)** microbiomes. Each node represents an ASV, annotated at the genus level, while edges indicate either positive (co-occurrence/green) or negative (mutual exclusion/red) correlations based on relative abundance profiles. Node size reflects the number of direct connections (Degree) among Taxa, whereas colours represent the betweenness centrality **(A-C)** and the modules **(B-D).**

## Discussion

In this study, we examined how localized microbiomes (ejaculate) and external microbiomes (skin) are linked to reproductive potential using male guppies (*Poecilia reticulata*). Comparisons of microbial diversity and community structure revealed clear niche differentiation. Skin microbiomes showed significantly higher alpha diversity compared to ejaculate-associated microbiota, as measured by total estimated richness (Chao1 index). This likely reflects the skin’s continuous exposure to the aquatic environment [42–44]. In contrast, the lower diversity of the ejaculate-associated microbiota, consistent with previous reports [45–48], reflects its development within immunologically protected internal compartments. In mammals, for example, testicular immune privilege is tightly regulated to prevent microbial intrusion [49], and similar principles may apply to fish, with tight host control over reproductive tract colonization limiting microbial variability.

Interestingly, the skin microbiome mirrored those in related species, such as *Gambusia affinis* [50,51], suggesting a conserved poeciliid core microbiota with dominant genera including *Acinetobacter*, *Sphingomonas*, *Massilia* and *Acidovorax*. In contrast, ejaculate-associated microbiomes showed distinct taxonomic profiles and lower inter-individual dispersion compared with skin microbiomes (figure 1A,B). We observed high inter-individual variability in microbial composition, with some samples dominated by specific genera, such as *Rheinheimera* and *Mycobacterium* (figure 1E). As all fish were housed under identical conditions, the observed variability highlights the influence of intrinsic host factors - such as social interactions and physiological state - on microbiome assembly. These factors, well documented in humans and other animals, can substantially shape microbiome structure and composition [52–55].

Our dbRDA analyses confirmed that tissue type is the dominant factor shaping bacterial community composition, explaining roughly 13–15% of the total constrained variance (p < 0.001). When sperm velocity (VCL) was added as a constraining variable, the combined model remained significant (adjusted R² = 0.125, p = 0.001), indicating that sperm performance explains a small but consistent portion of microbiome variation. However, when controlling for tissue identity, this relationship disappeared (adjusted R² ≈ 0.007, p ≈ 1), showing that the apparent VCL–microbiome association largely reflects broader between-tissue differences rather than within-tissue effects. This highlights that while microbial communities covary with sperm traits, the tissue environment exerts the primary structuring influence on microbiome composition. Genera such as *Mycobacterium, Massilia, Acinetobacter, Staphylococcus*, *Lactobacillus*, and *Pseudomonas* were among those most strongly associated with variation in sperm traits (Fig. 2B, and S3, tables S6 and S7).

Crucially, we found that bacterial richness in the skin microbiome was negatively associated with sperm velocity (VCL), which is a key predictor of fertilization success [29] (Fig 2A). No association was detected with ejaculate-associated microbiome diversity and VCL, consistent with findings on other species [56]. In humans, however, increased semen alpha diversity has been linked to higher sperm DNA fragmentation and infertility [16]. Our data show that males with higher-diversity skin communities exhibited slower sperm, suggesting that skin microbial diversity may reflect, or potentially influence, physiological states linked to reproductive performance. This negative correlation challenges the common assumption that greater diversity is inherently beneficial [44,57–61].

One hypothesis for the observed results is that compromised immunity in less healthy males leads to more diverse skin microbiota, as immune suppression is known to promote microbial overgrowth and compositional instability in fish[62–64]. The observed negative association between sperm velocity and skin microbial diversity may therefore represent a physiological trade-off between immune maintenance and reproductive investment. Conversely, healthier and more active males may sustain a lower-diversity but more stable skin microbial community, indicative of stronger host filtering and immune control. Minich et al. (2022) found that faster-swimming fish tended to have less diverse microbiota across body sites [65], suggesting that bolder males with better swimming endurance, and potentially better sperm performance [40], may maintain a more controlled skin microbiome [66–69].

While patterns of association are clear, causality remains unproven. The complexity of microbial interactions and the variability among individuals and taxa make it challenging to isolate causal pathways. Furthermore, the microbiomes of laboratory animals are known to differ markedly from those of their wild counterparts [70,71], meaning that specific associations or causal relationships present in natural populations may be diminished or lost in outbred laboratory stocks.

From an evolutionary standpoint, our findings are especially provocative. If traits like sperm velocity are influenced by microbial consortia beyond the reproductive tract, then sexual selection may act on host phenotypes, as well as on microbial structure, or on host traits that modulate it. This broadens the traditional view of sexual selection to include symbiotic organisms as integral components of the reproductive phenotype, emphasizing the importance of inter-organ crosstalk, particularly through microbiomes. In practical terms, the skin microbiome emerges as a promising non-invasive proxy for assessing reproductive health. If microbial diversity or taxon-specific signatures consistently correlate with sperm quality, they could serve as ethical, field-applicable tools for assessing fertility without disrupting the animal. Altogether, our results provide a foundation for investigating causal mechanisms and determining whether microbial consortia directly influence reproductive traits or serve as biomarkers of broader physiological conditions.

## Ethical statement

All experimental procedures were approved by the Animal Welfare Committee of the University of Padova and conducted in accordance with relevant national and EU legislation on the protection of animals used for scientific purposes (Directive 2010/63/EU). Permit number 57/2025-PR.

## Author’s contributions

M.G. conceived the study, and was responsible for data collection and curation, formal analysis, investigation, methodology, validation, visualization, and writing—original draft, review, and editing. G.L.G. contributed to data curation, formal analysis, methodology, validation, visualization, and writing—review and editing. A.D. contributed to data collection and curation, formal analysis, investigation, methodology, and writing—review and editing. L.P. contributed to data collection, validation, and writing—review and editing. A.Q. contributed to formal analysis, methodology, and validation. E.B. contributed to methodology, validation, and writing—review and editing. C.G. contributed to methodology, validation, writing—review and editing, and secured funding. M.F. contributed to validation, writing—review and editing, and secured funding. M.E.M. contributed to investigation, methodology, validation, writing—original draft, writing—review and editing, and secured funding.

All authors approved the final version of the manuscript.

## Data accessibility

All 16S rRNA gene sequencing data have been deposited to Genbank (see linked project PRJNA1284610). All data and code used in this study are provided in the electronic supplementary material.

## Acknowledgements

The authors thank Giulia Dalla Rovere and Rafaella Franch for technical support at the Department of Comparative Biomedicine and Food Science, University of Padova. We are also grateful to the technical staff of the Poeciliid rearing facilities at the Department of Biology, University of Padova.

