## Supplementary material for "From microbiome to sperm motility traits: An inside out perspective"

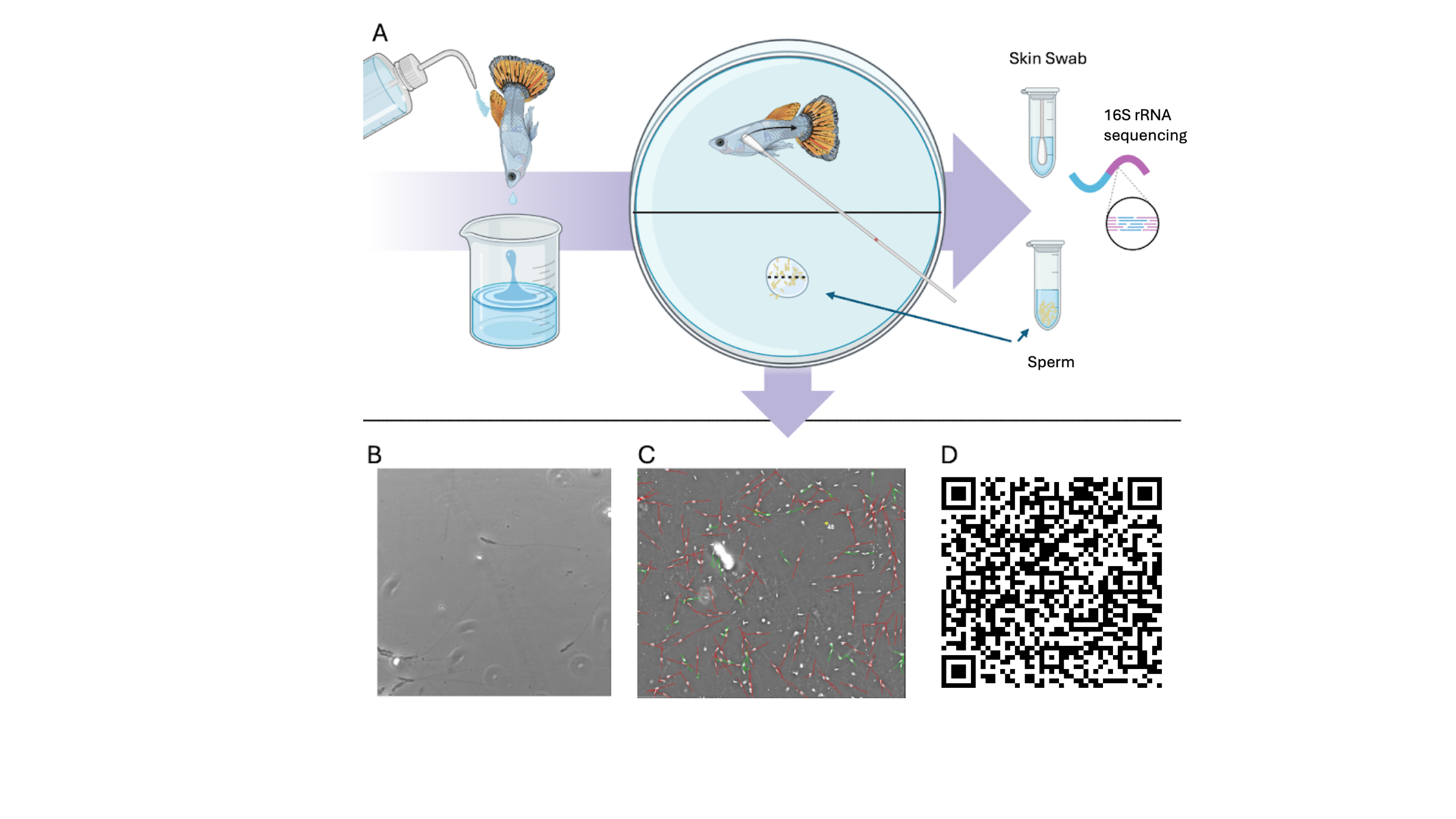

**Figure S1** **Overview of the experimental procedures.** **(A)** Sampling methodology, including collection of skin swabs and sperm bundles from Poecilia reticulata males, followed by 16S rRNA sequencing **(B)** Microscopic view of unprocessed sperm cells. **(C)** Sperm trajectories and velocities obtained using Computer-Assisted Sperm Analysis (CASA) software. **(D)** QR code linking to video showing swimming sperm cells with overlaid trajectory tracking.

***Extended methodology***

***SP1. Sampling and analysis contamination control procedures***

All microbiological sampling was conducted under strict aseptic conditions to prevent any cross-contamination between tissue types or individuals. Sterile gloves, face masks, and laboratory coats were worn throughout all procedures, and gloves were changed between individual fish and between tissue types. Each fish was anesthetized and processed individually on sterile Petri dishes. All working surfaces and instruments were thoroughly disinfected with 70% ethanol and RNaseZap™ spray before and between procedures. Sterile tweezers and scalpels were used and replaced between fish and tissue types.

Skin swabs were always collected first to ensure that subsequent sperm sampling could not introduce external contaminants. Following the skin collection, sperm bundles were obtained directly from the gonopore by gentle abdominal pressure onto a new sterile Petri dish containing physiological saline, ensuring that the sample did not contact the fish’s skin. All consumables (swabs, Petri dishes, microtubes) were sterile and opened immediately prior to use.

All tissue processing steps, including RNA extractions, were performed under a certified microbiological flume hood to maintain sterility and prevent environmental contamination. These precautions, combined with systematic surface decontamination between individuals, minimized potential carry-over contamination and ensured that the recovered microbial communities accurately represented the tissue-specific microbiota

***SP2. Diversity Metrics and Community Composition***

Chao1 and Shannon index were estimated in *Qiime2* using the command *diversity alpha.* Weighted UniFrac distance matrix were also estimated. These metrics require a rooted phylogenetic tree relating the features to one another. The rooted phylogenetic tree was generated with the align-to-tree-mafft-fasttree pipeline from the q2-phylogeny plugin [1,2]*.*

A non-parametric ANOVA (Wilcoxon test) was performed using the command *diversity alpha-group-significance* to assess any significant differences between skin and sperm bundles. Alpha diversity indices value and p-value were graphed using *ggpubr*. The beta-diversity was assessed by a Principal Coordinate Analysis (PCoA) based on the UniFrac distance matrix. Significance among skin and sperm bundles was tested using permutational analysis of variance (PERMANOVA) [3] by the *adonis2* function with 9999 permutations. The *adonis2* function come from the *vegan* package[4]. Differential abundance analysis was performed using ALDEx2[5,6] and a Benjamini and Hochberg (BH) false discovery rate correction was applied to the results[7]. To explore the relationships between most abundant genera and sperm traits, the Spearman correlations were estimated using the *Hmisc* package.

As for bacterial network construction, the following topological features used to describe network properties: (i) connectivity, which is the number of links (i.e. edges) of a node to other nodes; (ii) clustering coefficient, a measure of interconnectivity in the neighbourhood of a node; (iii) path length, which is the average number of edges on the shortest path connecting any two nodes and (iv) betweenness centrality, which reflects the number of times a node plays a role as a connector along the shortest path between two other nodes.

***SP3. Sperm Trait Distributions and Principal Component Analysis***

To evaluate the distribution of sperm motility parameters, we assessed fits for normal, gamma, and lognormal distributions. The *fitdistrplus* package[8] was used to fit each distribution to the variable, and model fit was compared through density plots and Q-Q plots. This approach enabled identification of the most suitable distribution model for downstream analyses. To quantify the consistency of sperm kinematic measurements across the three replicate sperm-bundle groups collected from each male, we estimated repeatability (R) using the *rptR* package[9] (Gaussian LMMs with male ID as a random factor; 1,000 bootstraps and 1,000 permutations). Most traits showed **moderate and statistically significant repeatability**, including VCL (R = 0.38), VAP (R = 0.45), VSL (R = 0.47), BCF (R = 0.42), and LIN (R = 0.32). ALH showed low and non-significant repeatability (R = 0.14), and STR showed no detectable repeatable variance (R ≈ 0).

Given that the majority of traits exhibited meaningful within-male repeatability, and because only one microbiome richness value (one Chao1/Shannon per male per tissue) was available, **replicate measurements were averaged per male** prior to PCA and microbiome–trait modelling. This approach prevents pseudoreplication while preserving reliable between-male variation.

Correlation matrices were computed to test collinearity among sperm trait metrics[10,11]. The *GGally*[12] and *corrplot*[13] packages were used to visualize these correlations, with Pearson’s correlation coefficients calculated between variables. Briefly, a matrix of Pearson’s correlation coefficients was generated for all pairs of variables in the subset. Significance testing of correlations was conducted using the *cor.mtest* function to calculate p-values for each pairwise correlation. Variables with significant correlations were extracted, and scatter plots and regression smoothers were added to highlight trends. *ggpairs* plots were generated, indicating the significance of pairwise relationships within the dataset. Using this method, collinearity between sperm metrics were detected; specifically, strong correlations were found among velocity measures (VAP–VSL r = 0.99; VAP–VCL r = 0.97; VSL–VCL r = 0.94; all p < 0.001), and several directionality traits were also associated (e.g., STR–LIN r = 0.72, p < 0.001; ALH–STR r = –0.74, p < 0.001). Correlations were calculated using GGally[12] and corrplot[13].

Given the strong intercorrelations among velocity-related and directionality traits, a Principal Component Analysis (PCA) was applied to the highly repeatable and collinear sperm parameters in the dataset, which included VCL, VAP, VSL, LIN, ALH, BCF, and STR. To ensure comparable scales across variables, data were centred and scaled prior to analysis using the *prcomp* function. Loadings for each motility parameter on the principal components were calculated and visualized using the *factoextra* package[14], with eigenvalues indicating the proportion of variance explained by each component. All PCA plots and variable contributions were visualized using the *factoextra*[14] and *ggplot2* [15] packages. Eigenvalue scree plots were generated to illustrate the variance explained by each principal component. Biplots were produced to visualize relationships between motility parameters and microbial taxa within the principal component space, facilitating interpretation of key drivers of variance in the dataset. Model fit and diagnostics were evaluated via the *sjPlot* package, and estimates were visualized to interpret significant effects. Coefficient plots were generated for each model to visualize significant predictors. Separate prediction plots were created to illustrate interaction terms and provide insights into relationships between motility parameters, diversity indices and specific bacterial abundances.

***SP4. Modelling Microbiome–Sperm Trait Interactions***

To identify how sperm functional traits covary with host-associated microbiome diversity, for each response variable (VCL, PC1, PC2), we implemented an automated, multi-step model selection pipeline in R (v4.4.1) that compared alternative parameterizations of bacterial diversity and data scaling.

### ***1. Determination of model structure and error distribution***

We first evaluated the most appropriate **error family** for sperm trait data (VCL, PC1, PC2) by comparing normal, gamma, and log-normal distributions. Linear models, Linear Mixed-Effects Models (LMMs) and Generalized Linear Mixed-Effects Models (GLMMs) were fitted using *lme4* [16] and *glmmTMB*[17]*.* Variance components were obtained via ***lmerTest*** [18]***,*** and model diagnostics (residual normality, heteroscedasticity) were checked using ***DHARMa*** [19]. Significance of fixed effects was assessed via **Type III Wald χ²** and **F-tests** with Satterthwaite’s approximation. All mixed-effects models included **male** as a random intercept, reflecting the paired sampling design (each male contributed one skin and one sperm-bundle microbiome)

### ***2. Initial tissue-explicit models***

To test whether the relationship between bacterial richness and sperm performance differed between tissues, we first fit tissue-explicit models of the form
  Y ~ Tissue × Chao1 + Shannon,

using both **raw** and **within-tissue standardized (z-scored)** Chao1 and Shannon values. This formulation allowed us to assess whether the relationship between microbial richness and sperm velocity differed across host compartments. Model performance across scaling variants was compared using AICc, and the best-performing formulation was retained for subsequent steps. These models quantified tissue-specific slopes and global interactions between tissue identity (skin vs. sperm bundles) and microbial alpha diversity.

### ***3. Component-specific mixed effect models (ChaoSkin / ChaoBundles)***

To decompose tissue effects into explicit richness terms for each compartment, we next fit paired-sample mixed effect models with random intercepts for individual ID:
  Y ~ ChaoSkin + ChaoBundles + (1 | ID).

Models were fitted as **LMMs or GLMMs (via lme4 / glmmTMB), depending on the error structure selected in step 1**. Predictors were z-standardized. Competing formulations (raw vs. standardized values; inclusion/exclusion of Shannon), were compared via **ΔAICc** using ***MuMIn::dredge,*** which automatically generated and ranked all model subsets from the full candidate structure. Multicollinearity among predictors was evaluated using ***performance::check_collinearity,*** leading to the exclusion of redundant variables when VIF > 5.

### ***4. Automated model selection and comparison***

For each response variable, the full candidate model set (including LM, LMM, and GLMM formulations where appropriate) was ranked using AIC, AICc, and BIC. Residual dispersion, diagnostic plots, and simulated residuals were examined using ***DHARMa*** to ensure correct error structure and detect overdispersion or zero inflation.
The most parsimonious model was selected as the one with the lowest AIC while retaining biological interpretability. Model comparisons were additionally validated using **likelihood-ratio tests** (anova()), enabling direct evaluation of competing error families, scaling choices, and predictor combinations.

### ***5. Final model evaluation and visualization***

Fixed effects were summarized using ***sjPlot::tab_model,*** and 95 % confidence intervals for marginal effects were derived via ***ggeffects***.

Per-tissue slopes and interactions were extracted using ***emmeans::emtrends,*** with model-specific significance tested via Wald χ². Goodness of fit was quantified by ***marginal (R²m)*** *and* ***conditional (R²c*)** coefficients obtained through ***MuMIn::r.squaredGLMM.***

The same modeling pipeline—error-family testing, automated model selection, and diagnostic evaluation—was applied to VCL, PC1 and PC2 as responses.

***7. Summary of modelling rationale***

Across all analyses, we modelled **VCL, PC1,** and **PC2** as response variables and compared linear models, linear mixed-effects models, and GLMMs differing in their error structure, scaling, and richness parameterization. Where supported by AICc, models included a **random intercept for male ID** to account for paired skin–bundle samples; however, in some cases (notably PC1 and PC2), the best-supported models were **simple linear models** without a random effect, indicating minimal between-male variance (see electronic supplementary files on Dryad).

Final Fixed effects consisted of either
**(i)** a Tissue × Chao1 interaction, or
**(ii)** tissue-specific richness terms (ChaoSkin and ChaoBundles).

This framework allowed us to quantify microbiome–trait associations both **between tissues** (interaction models) and **within tissues** (component-specific models), while incorporating random effects where they improved model performance.

***SP5. Multivariate analyses for Trait-Microbiome Relationships***

To investigate how sperm functional traits relate to microbial community composition across host tissues, we applied a two-stage automated constrained ordination framework implemented in R (v4.4.2) using the *vegan* and *permute* packages.

***Stage 1 – Automated model selection***

An initial automated model sweep (sweep_and_pick_rda) was performed to identify optimal abundance and prevalence thresholds as well as data transformations for robust ordination modeling. We systematically evaluated combinations of prevalence thresholds (2–7 samples per genus) and mean relative abundance cut-offs (0, 0.001, 0.0025, and 0.005), with optional square-root and taxon down-weighting transformations.

Each combination was fitted using both Constrained Correspondence Analysis (CCA) and distance-based Redundancy Analysis (dbRDA) based on Bray–Curtis dissimilarities. All models were assessed with 9,999 permutation tests, blocked by individual fish ID (*permute::how*(blocks = ID)), to respect the pairing of bundle (B) and skin (S) samples.

Model performance was summarized by adjusted R² and permutation-derived p-values.

The automated sweep revealed that dbRDA consistently captured more variance than CCA, and that moderate filtering criteria (prevalence ≥ 5 samples, mean relative abundance ≥ 0.005) produced the most stable and interpretable results.

***Stage 2 – Final optimized models***

Based on these outcomes, the final analyses were restricted to the 25 most prevalent genera to focus on the dominant and most consistent microbial contributors.

Two ordination methods were compared: a CCA, using chi-square distances on relative abundances; and a dbRDA, implemented via capscale with Bray–Curtis distances; an

For each method, two model structures were tested: (1) pooled models, relating community composition to sperm trait scores and tissue identity (Y ~ PC1 + PC2 + tissue or Y ~ VCL + tissue), and (2) within-tissue (partial) models, testing whether sperm traits explained community variation independently of tissue (Y ~ PC1 + PC2 + Condition(tissue) or Y ~ VCL + Condition(tissue)). Permutation significance was evaluated with blocked resampling by fish ID. Adjusted R² values were obtained using RsquareAdj, and ordination significance was assessed via anova.cca.

Ordination biplots were constructed at scaling 2, with tissue-coded sample points and arrows representing the constrained sperm trait axes (PC1, PC2, VCL). The 25 most informative genera were annotated by vector magnitude, and their species scores, angles, and cosines relative to each sperm trait axis were extracted to identify the key taxa driving the observed structure. To determine whether sperm trait–microbiome associations persisted within each host compartment, the same ordination models were repeated separately for bundle (B) and skin (S) subsets using the same top 25 genera and modeling parameters.

***Extended results (PC1 and PC2)***

Skin microbiome diversity was significantly associated with sperm traits principla components deriving from CASA. Higher diversity correlated with increased PC1 scores (velocity component; β = 0.94, p = 0.03), which, due to negative loadings, indicates lower sperm velocities (Figure S2, Tab. S3, S4). Conversely, PC2 scores (directionality) were negatively associated with skin diversity, suggesting more diverse skin microbiota leads to straighter sperm movement. (β = –0.74, p = 0.022, Figure S2, Tab. S3, S4).

In the RDAs, Genera such as Mycobacterium, Massilia, Acinetobacter, Brevundimonas, and Cutibacterium displayed the highest vector lengths, indicating that they contribute most to ordination structure. Mycobacterium showed strong positive alignment with PC1 (cos = 0.861), whereas several genera (e.g., Micrococcus, Bacillus, Staphylococcus) showed strong negative alignment with PC2, suggesting differential associations with sperm trajectory stability (Figure S3).

***SP7. Packages***

All the analyses were performed using RStudio version 4.4.2 (RStudio Team, 2024) equipped with the *car*[20], *glmmTMB*[17], *readxl*[21], *lme4*[16], *lmerTest*[18], *DHARMa*[22], *lsmeans*[23], *merTools*[24], *dplyr*[25], *tidyverse*[26], *rstatix*[27], *ggpubR*[28], *arsenal*[29], *knitr [30]*and *survival*[31] packages to perform exploratory analysis, run the main models, perform post-hoc tests, and create output tables. Graphical figures were plotted using *ggplot2*[15] , *ggpubr*[28], *sjPlot*[32], *sjmisc*[33], and *qqplotr*[34].

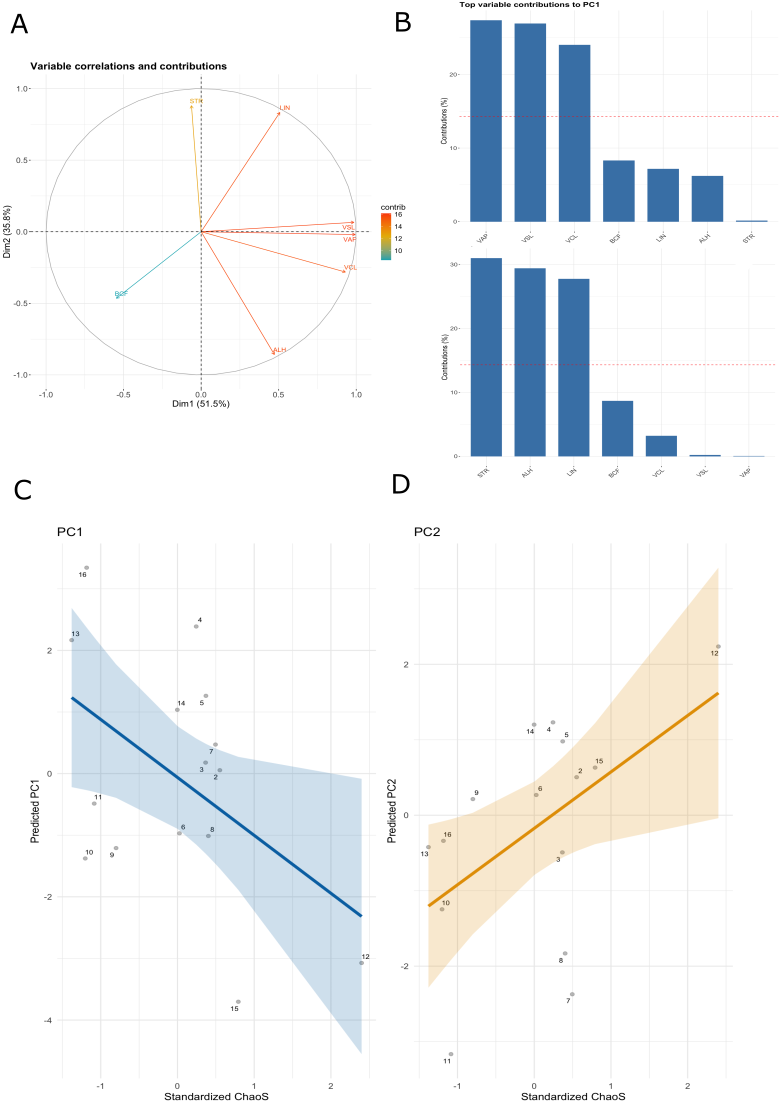

**Figure S2** **PCA explanatory variables** with their contribution to the components and correspondent directions **(A).** Single sperm motility variable contributions to dimension 1 and 2 (PC1, PC2**; (B)** and GLMM (glmmTMB) predictions showing the relationships between standardized ChaoS and the response variables: PC1 **(C)** and PC2 **(D)**, with 95% 204 confidence intervals. Individual data points are displayed with corresponding IDs.

**Table 1** **Detailed settings for CASA sperm motility acquisition and analysis.** Microscope setup, imaging frame rate, CASA software version, chamber preparation, sample temperature, activation solution, timing of video capture, analysis window, and motility thresholds used to classify static versus motile sperm are provided to ensure reproducibility of sperm kinematic measurements.

| **Parameter** | **Setting** |
| --- | --- |
| Microscope | Nikon Eclipse E200, phase-contrast |
| Objective | 10× phase-contrast |
| Camera frame rate | 50 fps |
| Video duration | 0.5 s per field |
| CASA system | Hamilton Thorne CEROS, version 12.3 |
| CASA calibration | Standard CEROS calibration for 10× phase contrast (pixel-to-µm based on microscope optical parameters) |
| Chamber type | Multitest slides, 12-well, MP coated with 1% polyvinyl alcohol solution |
| Sample/stage temperature | 25 ± 1 °C |
| Activation solution | 150 mM KCl + 2 mg mL⁻¹ BSA in deionized water. |
| Time post-activation | 0–5 min after activation depending on the sample |
| Analysis window | Continuous measurement during the first 5 min |
| Motility thresholds | Static cells defined as VAP < 20 μm s⁻¹ and VSL < 15 μm s⁻¹ |

**Table S2 Differential abundance of bacterial taxa between skin and sperm samples.** The table shows the average relative abundance (%) of each taxon in skin and sperm microbiomes, the mean difference between compartments, the associated p-values from paired statistical tests, and the adjusted p-values (p_adj) after multiple testing correction. Only taxa with non-zero mean abundance in at least one compartment are shown.

| **Bacterial Taxa** | **Average abundance (%)** | | **Mean_Diff** | **p_value** | **p_adj** |
| --- | --- | --- | --- | --- | --- |
|  | **Skin** | **Sperm** |  |  |  |
| 11-24 | 0.057 | 0.162 | -0.105 | 0.5516 | 0.6692 |
| 1174-901-12 | 0.379 | 0.046 | 0.333 | 0.3305 | 0.5004 |
| 28-YEA-48 | 0.027 | 0.171 | -0.144 | 0.0184 | 0.1401 |
| A4b | 0.324 | 0.324 | 0.000 | 0.1155 | 0.2736 |
| AKYG587 | 0.230 | 0.117 | 0.113 | 0.2208 | 0.3795 |
| AKYH767 | 0.153 | 0.188 | -0.035 | 0.6246 | 0.7157 |
| Acidibacter | 0.186 | 0.106 | 0.079 | 0.1051 | 0.2668 |
| Acidiphilium | 0.064 | 0.063 | 0.002 | 0.3533 | 0.5204 |
| Acidovorax | 0.163 | 0.034 | 0.129 | 0.0873 | 0.2442 |
| Acinetobacter | **0.424** | **2.976** | **-2.552** | **<0.001** | **0.0105** |
| Actinomyces | 0.031 | 0.172 | -0.141 | 0.1098 | 0.2705 |
| Aerococcus | 1.074 | 0.182 | 0.892 | 0.0745 | 0.2209 |
| Aeromonas | 0.115 | 0.062 | 0.053 | 0.9431 | 0.9665 |
| Akkermansia | 0.187 | 0.072 | 0.115 | 0.0400 | 0.1985 |
| Alloprevotella | 0.519 | 0.325 | 0.194 | 0.2075 | 0.3782 |
| Allorhizobium-Neorhizobium-Pararhizobium-Rhizobium | 0.062 | 0.492 | -0.430 | 0.1689 | 0.3482 |
| Anaerococcus | 0.164 | 0.284 | -0.121 | 0.1259 | 0.2807 |
| Aurantisolimonas | 0.084 | 0.015 | 0.069 | 0.5708 | 0.6824 |
| Bacillus | 0.307 | 0.526 | -0.218 | 0.8609 | 0.9047 |
| Bacteroides | 0.886 | 0.304 | 0.582 | 0.0187 | 0.1401 |
| Bdellovibrio | 0.119 | 0.156 | -0.037 | 1.0000 | 1.0000 |
| Bifidobacterium | 0.170 | 0.081 | 0.089 | 0.8677 | 0.9061 |
| Blastocatella | 0.137 | 0.113 | 0.024 | 0.6077 | 0.7062 |
| Blautia | 0.105 | 0.117 | -0.012 | 0.5154 | 0.6541 |
| Brevundimonas | **0.115** | **2.447** | **-2.332** | **<0.001** | **0.0105** |
| Bryobacter | 0.245 | 0.152 | 0.093 | 0.0803 | 0.2325 |
| Campylobacter | 0.179 | 0.070 | 0.109 | 0.0553 | 0.2068 |
| Candidatus_Kaiserbacteria | 0.522 | 0.368 | 0.154 | 0.2157 | 0.3782 |
| Candidatus_Solibacter | 0.225 | 0.105 | 0.120 | 0.0464 | 0.2004 |
| Cellvibrio | 0.050 | 0.143 | -0.094 | 0.8747 | 0.9077 |
| Chitinophaga | 0.040 | 0.126 | -0.086 | 1.0000 | 1.0000 |
| Christensenellaceae_R-7_group | 0.152 | 0.080 | 0.072 | 0.2178 | 0.3782 |
| Chryseobacterium | 0.033 | 0.180 | -0.147 | 0.0852 | 0.2424 |
| Cloacibacterium | 0.169 | 0.349 | -0.181 | 0.4186 | 0.5805 |
| Clostridia_UCG-014 | 1.436 | 0.445 | 0.991 | 0.0032 | 0.0521 |
| Clostridium_sensu_stricto_1 | 0.362 | 0.193 | 0.169 | 0.1256 | 0.2807 |
| Corynebacterium | 0.361 | 0.719 | -0.358 | 0.1839 | 0.3529 |
| Curvibacter | 0.054 | 0.212 | -0.158 | 0.0316 | 0.1800 |
| Cutibacterium | **0.647** | **2.732** | **-2.085** | **0.0020** | **0.0414** |
| Deinococcus | 0.521 | 0.806 | -0.285 | 0.3411 | 0.5070 |
| Dongia | 0.065 | 0.179 | -0.114 | 0.8315 | 0.8852 |
| Ellin6067 | 0.432 | 0.162 | 0.271 | 0.1218 | 0.2792 |
| Endobacter | 0.148 | 0.013 | 0.136 | 0.4498 | 0.6134 |
| Enhydrobacter | 1.378 | 2.478 | -1.100 | 0.0597 | 0.2082 |
| Enterococcus | 0.194 | 0.476 | -0.282 | 0.0120 | 0.1324 |
| Escherichia-Shigella | 0.691 | 0.506 | 0.186 | 0.5267 | 0.6623 |
| F082 | 0.128 | 0.115 | 0.013 | 0.5948 | 0.6960 |
| Flavobacterium | 0.295 | 0.365 | -0.070 | 0.5386 | 0.6633 |
| Fluviicola | 0.000 | 0.149 | -0.149 | 0.0050 | 0.0751 |
| Gemmatimonas | 0.120 | 0.085 | 0.035 | 0.2419 | 0.4031 |
| Gitt-GS-136 | 0.125 | 0.072 | 0.054 | 0.1145 | 0.2736 |
| HT002 | 0.149 | 0.072 | 0.077 | 0.0341 | 0.1873 |
| Haliangium | 0.227 | 0.129 | 0.098 | 0.0511 | 0.2057 |
| Halomonas | 0.142 | 0.059 | 0.083 | 0.9827 | 0.9955 |
| Hymenobacter | 0.112 | 0.155 | -0.043 | 0.2661 | 0.4348 |
| Inhella | 0.113 | 0.055 | 0.058 | 0.7816 | 0.8429 |
| JG30-KF-CM45 | 0.089 | 0.112 | -0.023 | 0.5338 | 0.6623 |
| JG30-KF-CM66 | 0.108 | 0.086 | 0.022 | 0.2092 | 0.3782 |
| KD4-96 | 0.378 | 0.306 | 0.072 | 0.2765 | 0.4430 |
| Klebsiella | 0.129 | 0.344 | -0.214 | 0.0120 | 0.1324 |
| Kocuria | 0.264 | 0.172 | 0.091 | 0.3947 | 0.5664 |
| Lachnospiraceae_NK4A136_group | 0.284 | 0.173 | 0.111 | 0.0741 | 0.2209 |
| Lachnospiraceae_XPB1014_group | 0.113 | 0.085 | 0.027 | 0.0427 | 0.1985 |
| Lactobacillales | **0.363** | **1.402** | **-1.038** | **<0.001** | **0.0105** |
| Lactobacillus | 2.356 | 1.619 | 0.737 | 0.1882 | 0.3529 |
| Latescibacteraceae | 0.121 | 0.067 | 0.054 | 0.2343 | 0.3967 |
| Latescibacterota | 0.420 | 0.291 | 0.129 | 0.4352 | 0.5984 |
| Lawsonella | 0.430 | 0.352 | 0.079 | 0.1604 | 0.3392 |
| Ligilactobacillus | 0.157 | 0.138 | 0.019 | 0.4067 | 0.5735 |
| Lysobacter | 0.212 | 0.082 | 0.131 | 0.0462 | 0.2004 |
| MND1 | 0.593 | 0.289 | 0.304 | 0.0433 | 0.1985 |
| Massilia | **0.348** | **3.924** | **-3.576** | **<0.001** | **0.0105** |
| Methylobacterium-Methylorubrum | **0.107** | **1.584** | **-1.477** | **<0.001** | **0.0184** |
| Methyloversatilis | 0.670 | 0.212 | 0.457 | 0.0965 | 0.2541 |
| Micrococcus | 0.657 | 0.633 | 0.024 | 0.0082 | 0.1042 |
| Moraxella | 0.019 | 0.148 | -0.129 | 0.0552 | 0.2068 |
| Morganella | **0.593** | **2.285** | **-1.692** | **<0.001** | **0.0184** |
| Muribaculaceae | 1.203 | 0.574 | 0.629 | 0.0644 | 0.2126 |
| Mycobacterium | 5.690 | 2.955 | 2.735 | 0.8916 | 0.9194 |
| NB1-j | 0.315 | 0.219 | 0.096 | 0.3095 | 0.4773 |
| NK4A214_group | 0.142 | 0.079 | 0.063 | 0.0970 | 0.2541 |
| Neochlamydia | 0.106 | 0.127 | -0.022 | 0.6246 | 0.7157 |
| Nevskia | 0.090 | 0.091 | 0.000 | 0.6550 | 0.7454 |
| Nitrospira | 0.468 | 0.257 | 0.211 | 0.0750 | 0.2209 |
| Niveispirillum | 0.041 | 0.177 | -0.136 | 0.5029 | 0.6432 |
| Nocardioides | 0.148 | 0.063 | 0.085 | 0.0952 | 0.2541 |
| Novosphingobium | **0.016** | **0.331** | **-0.315** | **<0.001** | **0.0184** |
| OM190 | 0.464 | 0.212 | 0.252 | 0.0611 | 0.2082 |
| OM27_clade | 0.074 | 0.108 | -0.034 | 0.7557 | 0.8258 |
| Obscuribacteraceae | 0.074 | 0.110 | -0.037 | 0.8240 | 0.8829 |
| Oceanobacillus | **0.375** | **1.243** | **-0.868** | **0.0023** | **0.0429** |
| Oscillospira | 0.262 | 0.080 | 0.181 | 0.0420 | 0.1985 |
| Other | 20.694 | 17.792 | 2.903 | 0.1751 | 0.3482 |
| PLTA13 | 0.202 | 0.119 | 0.084 | 0.1363 | 0.2998 |
| Paenibacillus | 0.108 | 0.191 | -0.083 | 0.1090 | 0.2705 |
| Pajaroellobacter | 0.040 | 0.100 | -0.060 | 0.2628 | 0.4336 |
| Parabacteroides | 0.152 | 0.067 | 0.085 | 0.0491 | 0.2026 |
| Paracoccus | 1.479 | 2.854 | -1.375 | 0.0267 | 0.1634 |
| Parcubacteria | 0.135 | 0.104 | 0.031 | 0.6775 | 0.7553 |
| Pedosphaeraceae | 0.230 | 0.112 | 0.118 | 0.0599 | 0.2082 |
| Perlucidibaca | 0.187 | 0.367 | -0.180 | 0.4149 | 0.5802 |
| Permianibacter | 0.088 | 0.135 | -0.046 | 0.7244 | 0.8022 |
| Phascolarctobacterium | 0.307 | 0.087 | 0.220 | 0.0164 | 0.1401 |
| Polaromonas | 1.154 | 0.024 | 1.131 | 0.6718 | 0.7553 |
| Pontimonas | 0.197 | 0.054 | 0.144 | 0.0180 | 0.1401 |
| Prevotella | 0.358 | 0.267 | 0.091 | 0.5891 | 0.6942 |
| Prevotella_7 | 1.158 | 0.031 | 1.127 | 0.1431 | 0.3105 |
| Prevotella_9 | 0.141 | 0.060 | 0.081 | 0.0727 | 0.2209 |
| Prevotellaceae_NK3B31_group | 2.386 | 0.574 | 1.812 | 0.0146 | 0.1401 |
| Prevotellaceae_UCG-001 | 0.421 | 0.112 | 0.309 | 0.0216 | 0.1483 |
| Prevotellaceae_UCG-003 | 0.172 | 0.106 | 0.066 | 0.2178 | 0.3782 |
| Proteus | 0.287 | 0.768 | -0.481 | 0.0366 | 0.1889 |
| Providencia | 0.100 | 0.392 | -0.292 | 0.0311 | 0.1800 |
| Pseudogracilibacillus | 0.063 | 0.215 | -0.152 | 0.3896 | 0.5639 |
| Pseudomonas | 2.568 | 1.309 | 1.259 | 0.4945 | 0.6420 |
| Psychrobacter | 0.159 | 0.102 | 0.057 | 0.3345 | 0.5018 |
| RB41 | 0.202 | 0.136 | 0.066 | 0.3144 | 0.4803 |
| Reyranella | 0.071 | 0.139 | -0.068 | 0.9835 | 0.9955 |
| Rheinheimera | 2.706 | 2.786 | -0.080 | 0.6744 | 0.7553 |
| Rhodococcus | 0.642 | 0.326 | 0.315 | 0.8556 | 0.9047 |
| Rikenellaceae_RC9_gut_group | 1.420 | 0.441 | 0.979 | 0.0130 | 0.1340 |
| Rokubacteriales | 0.857 | 0.336 | 0.521 | 0.0187 | 0.1401 |
| Romboutsia | 0.112 | 0.102 | 0.009 | 0.5428 | 0.6635 |
| Roseicyclus | 0.285 | 0.084 | 0.201 | 0.0258 | 0.1634 |
| Roseomonas | 0.030 | 0.172 | -0.141 | 0.1177 | 0.2736 |
| Rothia | 0.079 | 0.275 | -0.196 | 0.2845 | 0.4514 |
| Ruminococcus | 0.140 | 0.068 | 0.072 | 0.1177 | 0.2736 |
| S0134_terrestrial_group | 0.087 | 0.084 | 0.002 | 0.1777 | 0.3490 |
| SAR324_clade(Marine_group_B) | 0.138 | 0.137 | 0.000 | 0.3095 | 0.4773 |
| SBR1031 | 0.294 | 0.234 | 0.060 | 0.2765 | 0.4430 |
| SM1A02 | 0.259 | 0.263 | -0.004 | 0.4981 | 0.6420 |
| Saccharimonadales | 0.235 | 0.196 | 0.039 | 0.4565 | 0.6174 |
| Shewanella | 0.110 | 0.093 | 0.017 | 0.1733 | 0.3482 |
| Skermanella | 0.002 | 0.506 | -0.504 | 0.0062 | 0.0855 |
| Sphingomonas | 1.366 | 2.878 | -1.513 | 0.0717 | 0.2209 |
| Staphylococcus | 2.065 | 2.102 | -0.037 | 0.1015 | 0.2616 |
| Stenotrophomonas | 0.084 | 0.190 | -0.106 | 0.2157 | 0.3782 |
| Streptococcus | 1.397 | 1.121 | 0.277 | 0.2356 | 0.3967 |
| Streptomyces | 0.146 | 0.040 | 0.106 | 0.0739 | 0.2209 |
| Subgroup_10 | 0.160 | 0.087 | 0.073 | 0.0564 | 0.2068 |
| Subgroup_17 | 0.165 | 0.105 | 0.060 | 0.1449 | 0.3105 |
| Subgroup_2 | 0.312 | 0.214 | 0.098 | 0.3025 | 0.4753 |
| Subgroup_22 | 0.120 | 0.071 | 0.049 | 0.1872 | 0.3529 |
| TRA3-20 | 0.233 | 0.131 | 0.102 | 0.1630 | 0.3405 |
| Terrisporobacter | 0.123 | 0.091 | 0.033 | 0.3798 | 0.5545 |
| Thermoanaerobacterium | 0.105 | 0.021 | 0.083 | 0.5708 | 0.6824 |
| Timonella | 0.456 | 0.070 | 0.386 | 0.5793 | 0.6876 |
| Treponema | 0.952 | 0.304 | 0.649 | 0.0618 | 0.2082 |
| UCG-002 | 0.251 | 0.085 | 0.166 | 0.0258 | 0.1634 |
| UCG-005 | 0.250 | 0.099 | 0.151 | 0.0204 | 0.1463 |
| UCG-010 | 0.110 | 0.082 | 0.028 | 0.1871 | 0.3529 |
| Unclassified_Genus | 10.804 | 12.395 | -1.590 | 0.4945 | 0.6420 |
| Variovorax | 0.056 | 0.090 | -0.034 | 0.4981 | 0.6420 |
| Veillonella | 0.088 | 0.078 | 0.009 | 0.7617 | 0.8268 |
| Vibrio | 0.376 | 0.145 | 0.231 | 0.2017 | 0.3740 |
| Vicinamibacteraceae | 0.695 | 0.474 | 0.221 | 0.0902 | 0.2480 |
| WCHB1-41 | 0.140 | 0.052 | 0.089 | 0.0553 | 0.2068 |
| WD260 | 0.080 | 0.010 | 0.070 | 0.7526 | 0.8258 |
| [Eubacterium]_coprostanoligenes_group | 0.071 | 0.172 | -0.101 | 0.4724 | 0.6337 |
| bacteriap25 | 0.187 | 0.179 | 0.008 | 0.4039 | 0.5735 |
| dgA-11_gut_group | 0.359 | 0.131 | 0.229 | 0.0363 | 0.1889 |
| env.OPS_17 | 0.118 | 0.131 | -0.013 | 0.5338 | 0.6623 |
| p-251-o5 | 0.427 | 0.191 | 0.235 | 0.1719 | 0.3482 |
| p-2534-18B5_gut_group | 0.089 | 0.084 | 0.006 | 0.0474 | 0.2004 |
| uncultured | 4.479 | 3.638 | 0.841 | 0.4945 | 0.6420 |

**Table S3 Network topology and modularity metrics for skin and sperm bundles microbiome communities.**
The table summarizes key network statistics including node and link counts, number and effective number of modules, and modularity scores for both skin and sperm bundles microbial communities. Module composition details are reported, including node IDs and taxonomic affiliations assigned to each module. Taxa are grouped according to their primary module membership, highlighting structural community differences between body sites.

| **Skin Community** |  | **Sperm Community** |  |
| --- | --- | --- | --- |
| nodes on the original network: | 81 | nodes on the original network: | 56 |
| links on the original network: | 269 | links on the original network: | 135 |
| number of modules: | 4 | number of modules: | 3 |
| effective number of modules: | 2,02979507731575 | effective number of modules: | 1,23665029842283 |
| modularity | 0,446 | modularity | 0,487 |
| Detailed module info: |  | Detailed module info: |  |
| number of nodes | eff. number of nodes | number of nodes | eff. number of nodes |
| 65 | 52,658566 | 49 | 36,484215 |
| 8 | 7,896158 | 6 | 4,1528044 |
| 10 | 7,0688157 | 3 | 2,828427 |
| 5 | 3,807231 |  |  |
| node ID | the ID of the module where the node mostly belongs | node ID | the ID of the module where the node mostly belongs |
| 1174-901-12 | 1174-901-12 | Terrisporobacter | Candidatus_Kaiserbacteria |
| Acidiphilium | 1174-901-12 | Clostridium_sensu_stricto_1 | Candidatus_Kaiserbacteria |
| Aerococcus | 1174-901-12 | Candidatus_Kaiserbacteria | Candidatus_Kaiserbacteria |
| Endobacter | 1174-901-12 | Proteus | OM190 |
| Kocuria | 1174-901-12 | Oceanobacillus | OM190 |
| Thermoanaerobacterium | 1174-901-12 | Morganella | OM190 |
| Timonella | 1174-901-12 | Lactobacillales | OM190 |
| WD260 | 1174-901-12 | OM190 | OM190 |
| Bacteroides | Escherichia-Shigella | p-251-o5 | Prevotellaceae_NK3B31_group |
| Escherichia-Shigella | Escherichia-Shigella | WCHB1-41 | Prevotellaceae_NK3B31_group |
| Enhydrobacter | Escherichia-Shigella | AKYH767 | Prevotellaceae_NK3B31_group |
| Lachnospiraceae_NK4A136_group | Escherichia-Shigella | UCG-010 | Prevotellaceae_NK3B31_group |
| Muribaculaceae | Escherichia-Shigella | UCG-005 | Prevotellaceae_NK3B31_group |
| Lysobacter | Escherichia-Shigella | UCG-002 | Prevotellaceae_NK3B31_group |
| Prevotellaceae_UCG-001 | Escherichia-Shigella | Muribaculaceae | Prevotellaceae_NK3B31_group |
| Hymenobacter | Rheinheimera | Treponema | Prevotellaceae_NK3B31_group |
| Rheinheimera | Rheinheimera | Subgroup_22 | Prevotellaceae_NK3B31_group |
| Methyloversatilis | Rheinheimera | Sphingomonas | Prevotellaceae_NK3B31_group |
| Sphingomonas | Rheinheimera | Romboutsia | Prevotellaceae_NK3B31_group |
| AKYH767 | Vibrio | Paracoccus | Prevotellaceae_NK3B31_group |
| Alloprevotella | Vibrio | SBR1031 | Prevotellaceae_NK3B31_group |
| Candidatus_Kaiserbacteria | Vibrio | SAR324_clade(Marine_group_B) | Prevotellaceae_NK3B31_group |
| A4b | Vibrio | Saccharimonadales | Prevotellaceae_NK3B31_group |
| Christensenellaceae_R-7_group | Vibrio | Ruminococcus | Prevotellaceae_NK3B31_group |
| [Eubacterium]_coprostanoligenes_group | Vibrio | Akkermansia | Prevotellaceae_NK3B31_group |
| Clostridium_sensu_stricto_1 | Vibrio | Roseicyclus | Prevotellaceae_NK3B31_group |
| dgA-11_gut_group | Vibrio | Rokubacteriales | Prevotellaceae_NK3B31_group |
| F082 | Vibrio | Rikenellaceae_RC9_gut_group | Prevotellaceae_NK3B31_group |
| Flavobacterium | Vibrio | Prevotellaceae_UCG-003 | Prevotellaceae_NK3B31_group |
| bacteriap25 | Vibrio | Prevotellaceae_UCG-001 | Prevotellaceae_NK3B31_group |
| Gitt-GS-136 | Vibrio | Prevotellaceae_NK3B31_group | Prevotellaceae_NK3B31_group |
| Haliangium | Vibrio | Prevotella_9 | Prevotellaceae_NK3B31_group |
| JG30-KF-CM45 | Vibrio | Pontimonas | Prevotellaceae_NK3B31_group |
| JG30-KF-CM66 | Vibrio | PLTA13 | Prevotellaceae_NK3B31_group |
| Candidatus_Solibacter | Vibrio | Phascolarctobacterium | Prevotellaceae_NK3B31_group |
| Lactobacillus | Vibrio | Pedosphaeraceae | Prevotellaceae_NK3B31_group |
| Clostridia_UCG-014 | Vibrio | Parcubacteria | Prevotellaceae_NK3B31_group |
| Latescibacteraceae | Vibrio | Parabacteroides | Prevotellaceae_NK3B31_group |
| Campylobacter | Vibrio | dgA-11_gut_group | Prevotellaceae_NK3B31_group |
| Latescibacterota | Vibrio | Campylobacter | Prevotellaceae_NK3B31_group |
| MND1 | Vibrio | Oscillospira | Prevotellaceae_NK3B31_group |
| KD4-96 | Vibrio | Methylobacterium-Methylorubrum | Prevotellaceae_NK3B31_group |
| NB1-j | Vibrio | Massilia | Prevotellaceae_NK3B31_group |
| Nitrospira | Vibrio | Ligilactobacillus | Prevotellaceae_NK3B31_group |
| NK4A214_group | Vibrio | Lachnospiraceae_NK4A136_group | Prevotellaceae_NK3B31_group |
| OM190 | Vibrio | JG30-KF-CM66 | Prevotellaceae_NK3B31_group |
| Neochlamydia | Vibrio | F082 | Prevotellaceae_NK3B31_group |
| OM27_clade | Vibrio | Gitt-GS-136 | Prevotellaceae_NK3B31_group |
| Oscillospira | Vibrio | AKYG587 | Prevotellaceae_NK3B31_group |
| p-251-o5 | Vibrio | Clostridia_UCG-014 | Prevotellaceae_NK3B31_group |
| p-2534-18B5_gut_group | Vibrio | Christensenellaceae_R-7_group | Prevotellaceae_NK3B31_group |
| Parabacteroides | Vibrio | Candidatus_Solibacter | Prevotellaceae_NK3B31_group |
| Parcubacteria | Vibrio | Brevundimonas | Prevotellaceae_NK3B31_group |
| Pedosphaeraceae | Vibrio | Acinetobacter | Prevotellaceae_NK3B31_group |
| Phascolarctobacterium | Vibrio | Bacteroides | Prevotellaceae_NK3B31_group |
| PLTA13 | Vibrio | Alloprevotella | Prevotellaceae_NK3B31_group |
| Prevotella | Vibrio |  |  |
| Prevotella_9 | Vibrio |  |  |
| Prevotellaceae_NK3B31_group | Vibrio |  |  |
| Prevotellaceae_UCG-003 | Vibrio |  |  |
| Psychrobacter | Vibrio |  |  |
| Rikenellaceae_RC9_gut_group | Vibrio |  |  |
| Rokubacteriales | Vibrio |  |  |
| AKYG587 | Vibrio |  |  |
| Roseicyclus | Vibrio |  |  |
| Ruminococcus | Vibrio |  |  |
| Saccharimonadales | Vibrio |  |  |
| SAR324_clade(Marine_group_B) | Vibrio |  |  |
| SBR1031 | Vibrio |  |  |
| Subgroup_17 | Vibrio |  |  |
| Subgroup_2 | Vibrio |  |  |
| Terrisporobacter | Vibrio |  |  |
| TRA3-20 | Vibrio |  |  |
| Treponema | Vibrio |  |  |
| UCG-002 | Vibrio |  |  |
| UCG-005 | Vibrio |  |  |
| uncultured | Vibrio |  |  |
| Variovorax | Vibrio |  |  |
| Vibrio | Vibrio |  |  |
| Vicinamibacteraceae | Vibrio |  |  |
| WCHB1-41 | Vibrio |  |  |

### ***Table S4 Across-tissues models testing the association between sperm motility traits (VCL, PC1, PC2) and bacterial richness (Chao₁) across tissues.*** Estimates (β) indicate effect size; SE denotes standard error; t/z values refer to test statistics from model summaries. *As the best-supported models (based on AICc) were simple linear models without random effects, marginal and conditional R² are identical; therefore, only adjusted R² (AdjR²) is reported. Statistically significant effects (p < 0.05) appear in* ***bold*.**

| **Response** | **Term** | **Estimate (β)** | **SE** | **t / z value** | **p value** | **AdjR²** |
| --- | --- | --- | --- | --- | --- | --- |
| **VCL** | (Intercept) | 120.000 | 7.660 | 15.70 | **<0.001** | 0.103 |
|  | **Chao1** | **–0.0089** | **0.0036** | **–2.46** | **0.021** |  |
|  | Tissue (B) | –13.700 | 9.700 | –1.41 | 0.170 |  |
|  | Chao1 × Tissue (B) | 0.0057 | 0.0070 | 0.81 | 0.425 |  |
| **PC1** | (Intercept) | 1.560 | 0.900 | 1.73 | 0.095 | 0.051 |
|  | **Chao1** | **–0.00087** | **0.00043** | **–2.04** | **0.052** |  |
|  | Tissue (B) | –1.23 | 1.14 | –1.08 | 0.292 |  |
|  | Chao1 × Tissue (B) | 0.00041 | 0.00083 | 0.50 | 0.624 |  |
| **PC2** | (Intercept) | –1.04 | 0.74 | –1.40 | 0.174 | 0.003 |
|  | Chao1 | 0.00057 | 0.00035 | 1.65 | 0.112 |  |
|  | Tissue (B) | 0.85 | 0.94 | 0.90 | 0.375 |  |
|  | Chao1 × Tissue (B) | –0.00032 | 0.00068 | –0.47 | 0.646 |  |

### ***Table S5 Within-tissue mixed-effect models summaries testing the association between skin microbial richness (Chao₁ₛ) and sperm motility traits (VCL, PC1, PC2).*** *Estimates (β) indicate effect size; SE denotes standard error; t/z values refer to test statistics from model summaries. R²ₘ represents the marginal R² (variance explained by fixed effects only), and R²c the conditional R² (including random effects). Significant effects (p < 0.05) are shown in bold.*

| **Response** | **Term** | **Estimate (β)** | **SE** | **t / z value** | **p value** | **R²ₘ** | **R²c** |
| --- | --- | --- | --- | --- | --- | --- | --- |
| **VCL** | (Intercept) | 104.107 | 3.319 | 31.366 | **< 0.001** | 0.391 | 0.990 |
|  | Chao1ₛ | –10.305 | 3.435 | –3.00 | **0.002** |  |  |
| **PC1** | (Intercept) | 0.062 | 0.425 | 0.145 | 0.884 | 0.245 | 0.622 |
|  | Chao1ₛ | 0.940 | 0.440 | 2.13 | **0.032** |  |  |
| **PC2** | (Intercept) | 0.174 | 0.316 | 0.550 | 0.582 | 0.271 | 0.635 |
|  | Chao1ₛ | –0.747 | 0.327 | –2.28 | **0.022** |  |  |

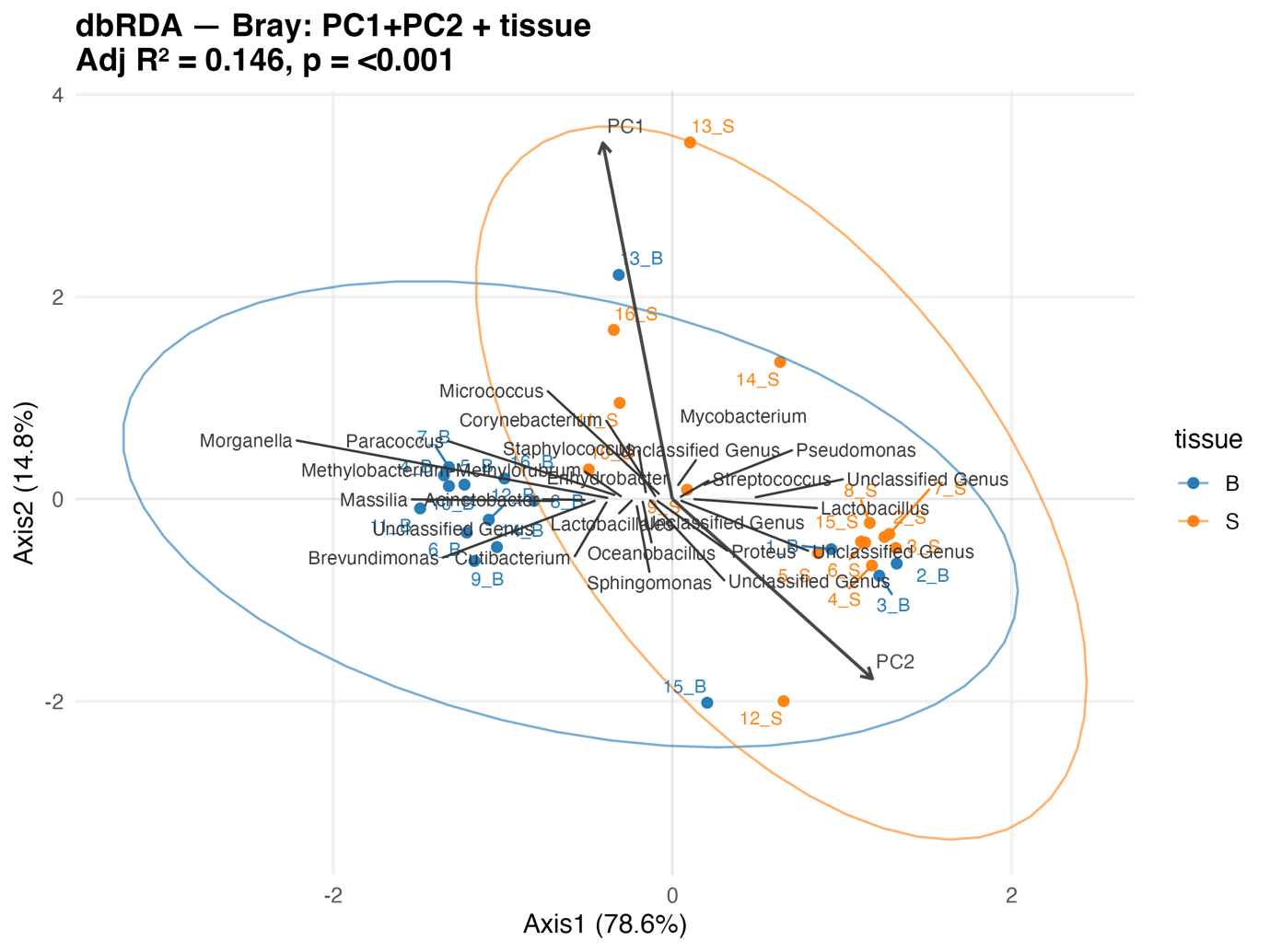

***Figure S3*** ***Distance-based redundancy analysis (dbRDA, Bray–Curtis) showing bacterial community composition constrained by PC1, PC2 and tissue type****. Ellipses (95% confidence) illustrate tissue-level clustering. PC1 and PC2 vectors indicate the direction of increasing sperm principal components and their association with specific genera.*

**Table S6 *Key microbial taxa contributing to variation in community structure along the VCL-constrained RDA axis.***
Values represent the coordinates of each genus vector on the first constrained axis (**sx**) and the secondary unconstrained axis (**sy**). **Vector length** corresponds to the magnitude of each taxon’s contribution to the ordination space, while **angle_deg** indicates the orientation of the vector in degrees. The **cos_VCL** value represents the cosine of the angle between the taxon vector and the VCL constraint axis, providing an index of alignment with the sperm-velocity gradient. Values close to **1** indicate strong **positive** alignment; values close to **–1** indicate strong **negative** alignment; values near **0** suggest weak or no association.

| **Taxa** | **sx** | **sy** | **length** | **angle_deg** | **cos_VCL** |
| --- | --- | --- | --- | --- | --- |
| **Massilia** | -0.718 | -0.024 | 0.718 | -178.107 | 0.136 |
| **Mycobacterium** | 0.328 | 0.611 | 0.694 | 61.745 | 0.788 |
| **Acinetobacter** | -0.507 | -0.014 | 0.507 | -178.440 | 0.142 |
| **Unclassified Genus** | 0.486 | 0.016 | 0.486 | 1.855 | -0.137 |
| **Brevundimonas** | -0.466 | -0.010 | 0.466 | -178.789 | 0.148 |
| **Cutibacterium** | -0.420 | -0.009 | 0.420 | -178.720 | 0.146 |
| **Morganella** | -0.347 | 0.003 | 0.347 | 179.512 | 0.177 |
| **Sphingomonas** | -0.303 | -0.047 | 0.306 | -171.203 | 0.016 |
| **Paracoccus** | -0.302 | 0.045 | 0.305 | 171.507 | 0.312 |
| **Methylobacterium-Methylorubrum** | -0.301 | 0.012 | 0.301 | 177.708 | 0.208 |
| **Enhydrobacter** | -0.247 | 0.051 | 0.252 | 168.433 | 0.363 |
| **Pseudomonas** | 0.202 | 0.149 | 0.251 | 36.401 | 0.449 |
| **Lactobacillales** | -0.210 | -0.002 | 0.210 | -179.443 | 0.159 |
| **Oceanobacillus** | -0.175 | -0.007 | 0.176 | -177.877 | 0.132 |
| **Unclassified Genus** | 0.143 | -0.079 | 0.163 | -28.884 | -0.624 |
| **Unclassified Genus** | -0.001 | 0.139 | 0.139 | 90.233 | 0.986 |
| **Unclassified Genus** | -0.134 | 0.003 | 0.134 | 178.743 | 0.190 |
| **Lactobacillus** | 0.125 | -0.001 | 0.125 | -0.526 | -0.178 |
| **Proteus** | -0.099 | -0.003 | 0.099 | -178.479 | 0.142 |
| **Corynebacterium** | -0.072 | 0.001 | 0.072 | 178.833 | 0.189 |
| **Bacillus** | -0.056 | 0.040 | 0.069 | 144.144 | 0.714 |
| **Unclassified Genus** | -0.062 | -0.004 | 0.062 | -176.632 | 0.110 |
| **Staphylococcus** | -0.034 | 0.049 | 0.060 | 124.707 | 0.906 |
| **Streptococcus** | 0.042 | 0.011 | 0.043 | 14.744 | 0.088 |
| **Micrococcus** | -0.010 | 0.037 | 0.038 | 105.702 | 0.995 |

**Table S7 *Taxa most strongly associated with sperm motility gradients represented by PC1 (velocity component) and PC2 (directionality component), as identified by constrained RDA.*** Columns **sx**, **sy**, **length**, and **angle_deg** describe the genus loadings in RDA space. **cos_PC1** and **cos_PC2** are the cosines of the angles between the taxon vector and the PC1 or PC2 constrained axes, respectively. High absolute cosine values indicate strong alignment with that motility component.

| **Taxa** | **sx** | **sy** | **length** | **angle_deg** | **cos_PC1** | **cos_PC2** |
| --- | --- | --- | --- | --- | --- | --- |
| **Mycobacterium** | 0.306 | 0.685 | 0.750 | 65.929 | 0.861 | -0.494 |
| **Massilia** | -0.748 | -0.025 | 0.749 | -178.086 | 0.081 | -0.565 |
| **Acinetobacter** | -0.551 | -0.013 | 0.551 | -178.624 | 0.090 | -0.573 |
| **Unclassified Genus** | 0.522 | 0.016 | 0.522 | 1.745 | -0.084 | 0.567 |
| **Brevundimonas** | -0.486 | -0.013 | 0.486 | -178.488 | 0.088 | -0.571 |
| **Cutibacterium** | -0.417 | -0.017 | 0.417 | -177.612 | 0.073 | -0.558 |
| **Morganella** | -0.407 | 0.005 | 0.407 | 179.291 | 0.126 | -0.602 |
| **Paracoccus** | -0.363 | 0.032 | 0.365 | 174.969 | 0.201 | -0.660 |
| **Methylobacterium-Methylorubrum** | -0.321 | 0.012 | 0.321 | 177.913 | 0.150 | -0.621 |
| **Lactobacillales** | -0.255 | -0.000 | 0.255 | -179.925 | 0.113 | -0.591 |
| **Enhydrobacter** | -0.241 | 0.028 | 0.243 | 173.465 | 0.226 | -0.680 |
| **Pseudomonas** | 0.186 | 0.138 | 0.231 | 36.563 | 0.500 | -0.004 |
| **Sphingomonas** | -0.223 | -0.054 | 0.229 | -166.479 | -0.121 | -0.387 |
| **Oceanobacillus** | -0.196 | -0.008 | 0.196 | -177.737 | 0.075 | -0.560 |
| **Staphylococcus** | -0.185 | 0.049 | 0.192 | 165.245 | 0.363 | -0.778 |
| **Unclassified Genus** | 0.176 | -0.066 | 0.188 | -20.641 | -0.457 | 0.838 |
| **Unclassified Genus** | -0.156 | 0.003 | 0.156 | 178.728 | 0.136 | -0.610 |
| **Unclassified Genus** | 0.031 | 0.125 | 0.129 | 76.274 | 0.938 | -0.642 |
| **Lactobacillus** | 0.122 | 0.001 | 0.122 | 0.571 | -0.104 | 0.584 |
| **Proteus** | -0.110 | -0.006 | 0.111 | -176.993 | 0.062 | -0.549 |
| **Corynebacterium** | -0.106 | 0.010 | 0.106 | 174.545 | 0.208 | -0.666 |
| **Micrococcus** | -0.083 | 0.037 | 0.091 | 156.209 | 0.505 | -0.867 |
| **Unclassified Genus** | -0.078 | -0.006 | 0.079 | -175.974 | 0.044 | -0.534 |
| **Bacillus** | -0.061 | 0.032 | 0.069 | 152.070 | 0.566 | -0.901 |
| **Streptococcus** | 0.039 | 0.011 | 0.041 | 16.194 | 0.168 | 0.344 |
